# Development of a rocking bioreactor strategy to augment pro-angiogenic factor secretion by human adipose-derived stromal cells

**DOI:** 10.64898/2026.08.17.745211

**Authors:** Zhiyu Liang, Connor J. Gillis, Olga Trichtchenko, Tamie L. Poepping, Lauren E. Flynn

## Abstract

Cell therapies involving human adipose-derived stromal cells (hASCs) have shown promise for a range of clinical applications due to their ability to stimulate angiogenesis and dampen inflammation via paracrine mechanisms. However, a major barrier to the successful clinical translation of hASC-based therapies is that standard culture methods for expansion on rigid 2D tissue-culture polystyrene under static conditions diminish the pro-regenerative functionality of the cells. To address these limitations, the current project focused on the development of an *in vitro* bioreactor system for preconditioning hASCs to augment their capacity to stimulate regeneration through paracrine mechanisms. Specifically, the combined effects of decellularized adipose tissue (DAT) coatings, shear-stress stimulation, and varying oxygen tensions on hASC expansion and paracrine factor secretion were assessed. Additional studies were performed to characterize the effects of stimulating hASCs within the rocking bioreactor system using the pro-inflammatory cytokines IFN-γ and TNF-α. Expansion in the bioreactor under all conditions supported hASC growth with no observable morphological differences. However, dynamic culture on DAT coatings enhanced intracellular indoleamine 2,3-dioxygenase (IDO) expression in hASCs cultured under 20% O_2_. Moreover, culturing under dynamic conditions and/or on DAT coatings significantly increased secretion of the pro-angiogenic factors VEGF, HGF, and angiogenin. When pro-inflammatory cytokine priming was introduced, the expression of all tested paracrine factors was enhanced, particularly the immunomodulatory factors IL-6, IL-8 and MCP-1. Overall, a novel bioreactor system was developed for hASC expansion and preconditioning, demonstrating that the cell microenvironment can be tuned to modulate hASC paracrine factor secretion.

## 1 Introduction

Cell therapies involving human adipose-derived stromal cells (hASCs) have shown promise for a range of clinical applications due to their ability to produce paracrine factors that stimulate angiogenesis and dampen inflammation [1]. However, conventional methods for hASC expansion involving 2D culture on rigid tissue-culture plastic substrates can lead to cell senescence and an altered secretome that may limit their capacity to promote regeneration [2]. There is a clear need for improved cell manufacturing technologies to support the clinical translation of hASC-based therapies. Bioreactor platforms can enable large-scale cell expansion and can provide a means to precondition the cells by controlling their microenvironment to maintain or potentially enhance hASC pro-regenerative functionality [3].

A variety of preconditioning strategies have been investigated with the goal of modulating the hASC secretome, including cell-culture platforms that integrate biomaterial substrates [4–9], hypoxia [10,11], mechanical stimulation [12,13], and pro-inflammatory cytokine priming [14–16]. Most studies to date have focused on characterizing the effects of individual factors on the cellular response. For example, in the context of biomaterials, decellularized adipose tissue (DAT) scaffolds have shown promise as a tissue-specific, cell-instructive scaffold for ASC culture and delivery [17,18] that can support ASC proliferation [4,19], adipogenic differentiation [8,18,19], and *in vivo* angiogenesis [8]. Similarly, culture under reduced oxygen levels is a well-established mesenchymal stromal cell (MSC) preconditioning strategy that can stimulate pro-angiogenic paracrine activity through activation of hypoxia inducible factor-1α (HIF-1α) [20–22]. While less well-characterized, mechanical stimulation can also modulate MSC paracrine function. Exposure to shear stress has been shown to enhance pro-angiogenic factor secretion by hASCs [23,24], as well as modulate the ability of human MSCs to suppress TNF-α secretion by activated immune cells [25,26]. Finally, cytokine priming through exposure to pro-inflammatory factors such as IFN-γ and/or TNF-α has also been widely investigated as a strategy to enhance the immunosuppressive properties of MSCs [14,15]. For example, priming with IFN-γ and TNF-α has been shown to upregulate indoleamine 2,3-dioxygenase (IDO) expression in MSCs, augmenting their capacity to inhibit T-cell proliferation and promote M2-like macrophage polarization [27].

Recognizing the individual potential of each of these hASC preconditioning strategies, the current study sought to systematically characterize the effects of combining DAT coatings, hypoxia, and shear-stress stimulation on hASC growth and the secretion of pro-angiogenic and immunomodulatory paracrine factors. This combined approach is supported by previous studies applying a scaffold-based perfusion bioreactor [28,29]. More specifically, dynamic culturing of hASCs on DAT scaffolds within the perfusion bioreactor system under 2% O_2_ was shown to significantly enhance the hASC density and alter the hASC phenotype in the scaffold periphery relative to statically cultured controls [28,29]. In addition, dynamic culture on the DAT scaffolds under shear stress changed the levels of paracrine factors detected in scaffold lysates relative to the statically cultured controls, with significantly higher levels of pro-angiogenic HGF, pro-regenerative IL-10, and the macrophage chemoattractant CXCL10, along with significantly lower levels of pro-inflammatory IL-6 [29]. Importantly, *in vivo* testing in a subcutaneous implant model in athymic nude mice revealed there was a more pro-regenerative macrophage response, as well as enhanced angiogenesis and adipogenesis, in the DAT scaffolds that had been cultured within the perfusion bioreactor system under 2% O_2_ relative to statically cultured, freshly seeded, and unseeded controls [8,29].

While this preclinical data was promising, a critical limitation was that the perfusion bioreactor would be very challenging to scale-up, which would be required for clinical applications in humans. In addition, a heterogeneous hASC phenotype was observed within the DAT scaffolds, likely because the cells in the scaffold interior were subjected to varying microenvironmental cues – including varying levels of shear stress – compared to the cells in the scaffold periphery [28,29]. With the goal of developing a more clinically relevant and scalable approach, a rocking-bed bioreactor platform was developed in the current study to enable more controlled shear-stress stimulation of the hASCs. Using this new rocking bioreactor, we characterized the combined effects of DAT coatings and shear-stress stimulation on hASC expansion and paracrine factor secretion under varying oxygen tensions (2% versus 20% O_2_) with the goal of identifying microenvironmental conditions that would support hASC growth and augment hASC secretion of pro-angiogenic and immunomodulatory factors. In addition, we performed follow-up studies to characterize the effects of further stimulating the hASCs within the rocking bioreactor system with the pro-inflammatory cytokines IFN-γ and TNF-α on IDO expression and paracrine factor secretion.

## Materials and Methods

### 2.1. Materials

Unless otherwise stated, all chemicals and reagents were purchased from Sigma Aldrich Canada Ltd. (Oakville, Canada), and all antibodies were purchased from Abcam (Cambridge, UK).

### 2.2. Adipose tissue collection and processing

Subcutaneous adipose tissue samples were collected with informed consent from female patients undergoing elective breast or abdominal reduction surgeries at the London Health Sciences Centre (London, ON, Canada) with human research ethics board approval from Western University (HSREB 105426). The adipose tissue samples were processed within 2 h, following published protocols for hASC isolation [30] or decellularization [18]. Isolated hASCs were cultured on tissue-culture polystyrene (TCPS; Corning, NY, USA) at 37°C and 5% CO_2_ in complete proliferation media composed of DMEM/F12 supplemented with 10% fetal bovine serum (FBS), 100 U/mL penicillin, and 0.1 mg/mL streptomycin. hASCs at passage 3 or 4 were used for all studies.

#### 2.2.1. DAT coating fabrication

Decellularized tissue samples from 5-6 donors were pooled and processed following published protocols to generate DAT suspensions (25 mg/mL) in 0.2 M acetic acid [19]. To produce the coatings, the DAT suspension was applied to the wells of 8-well rectangular plates (Nunc, ThermoFisher) at 125 μL/cm^2^. For immunohistochemical analyses, square glass coverslips were placed in the centre of the wells prior to applying the coatings. The coatings were left in a biological safety cabinet to dry overnight. For disinfection in preparation for culture, the DAT coatings were incubated three times for 30 min each in sterile 70% ethanol, followed by one 30-min incubation in sterile 35% ethanol. The coatings were then rinsed three times using sterile PBS before seeding.

### 2.3. Bioreactor setup and ASC seeding

The rocking bioreactor consisted of the 8-well rectangular plates as cell-culture vessels, a compact digital rocker (Ohaus) to introduce shear stress, and a HypOxystation H35 system (HypOxygen) to maintain the hypoxic environment (2% O_2_). A standard cell-culture incubator (37°C, 95% air/5% CO_2_) was used for the normoxic (∼20% O_2_) culture conditions. Figure 1 provides an experimental overview and highlights the system setup, including the placement of the cell-culture plates on the rocker. Only four wells within the plates were used for cell culture to maintain a consistent axis of rotation across all samples. The remaining wells were filled with sterile PBS to reduce media evaporation.

**Figure 1.**
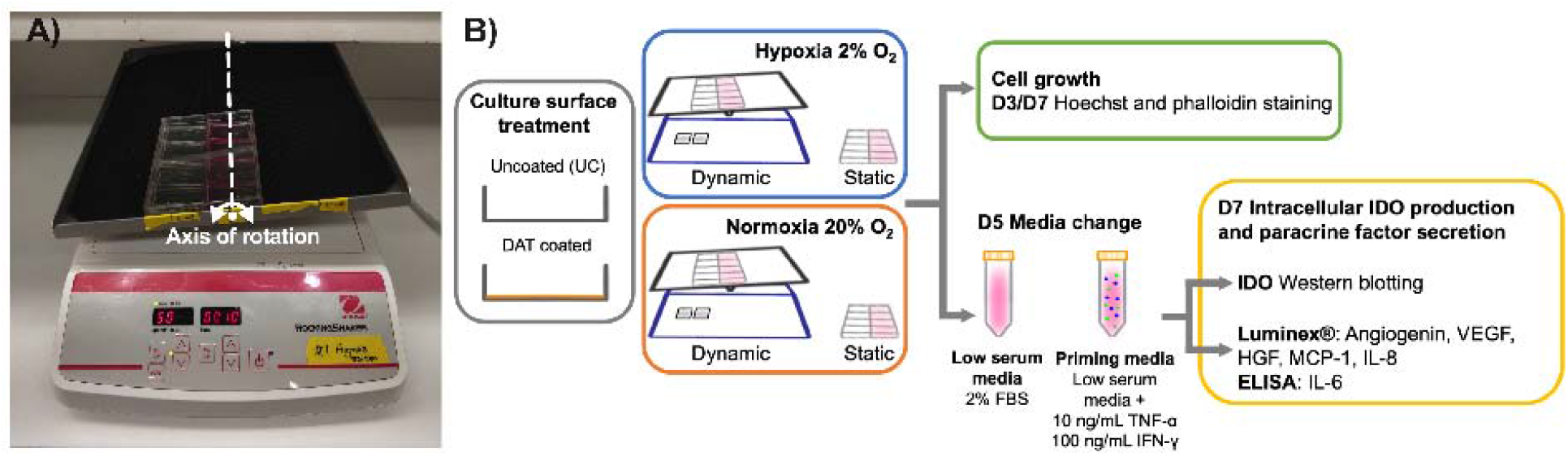
Experimental methods overview. A) Setup of the rocking bioreactor. The placement of the culture plate ensured that the centres of the wells in use were aligned with the axis of rotation of the rocker. B) Overview of the experimental methods.

**Figure 1.**
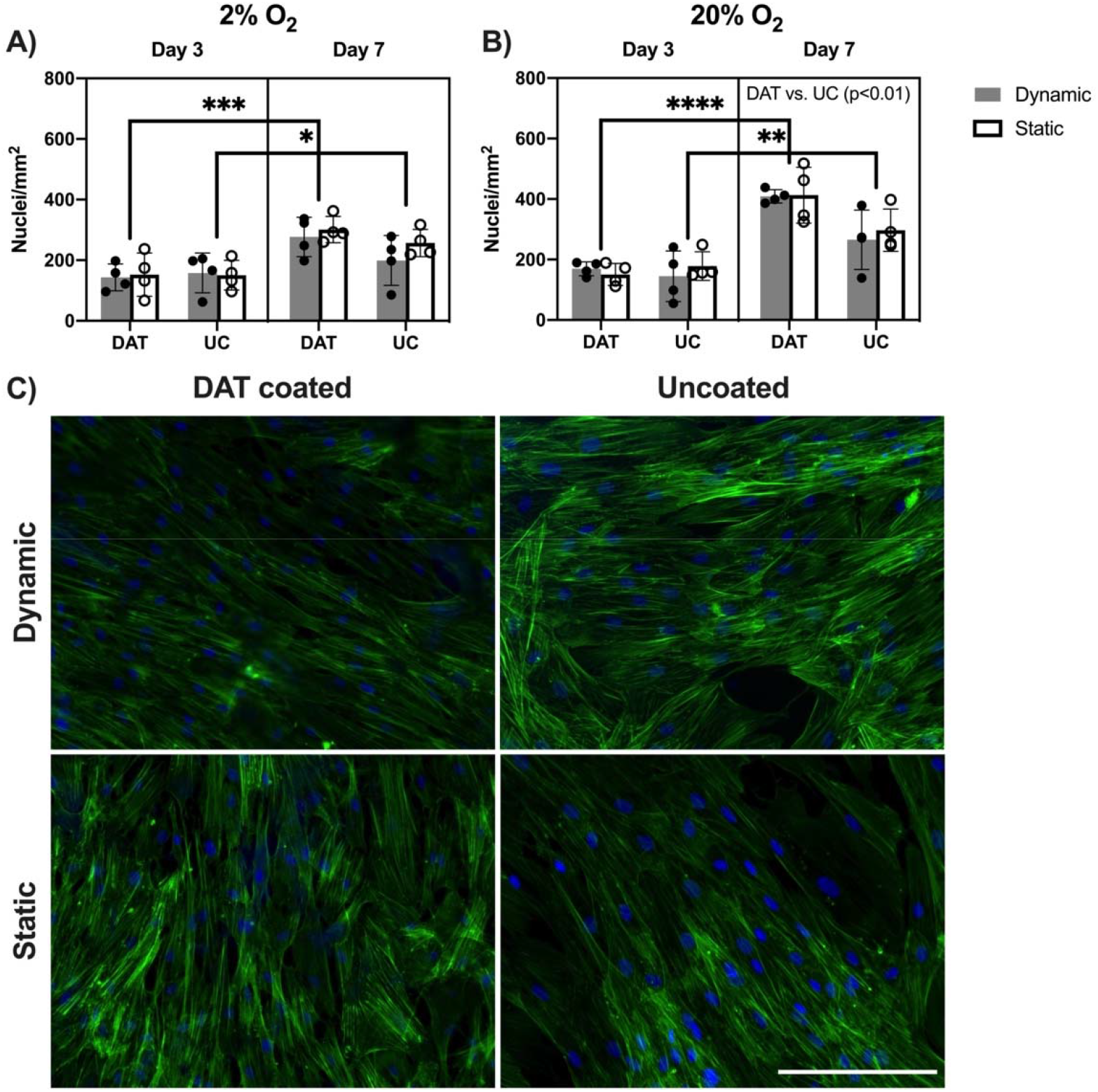
Dynamic culture with the rocking condition of 11° and 50 rpm did not substantially alter the hASC density at 7 days. Quantification of cell density after 3 and 7 days of culture under A) 2% O_2_ or B) 20% O_2_. Main effects analysis showed that cell density in the 20% O_2_ group was significantly higher than the 2% O_2_ group (p < 0.05). In addition, based on main effects analysis, the cell density in the DAT-coated group was significantly higher than the uncoated group under 20% O_2_ (p < 0.01). Based on two-way ANOVA analysis, the density of hASCs cultured on DAT coatings (***p < 0.001) and uncoated TCPS (*p < 0.05) under 2% O_2_ was significantly higher at 7 days as compared to 3 days. Similarly, under 20% O_2_, the density of hASCs cultured on DAT coatings (****p < 0.0001) and TCPS (**p < 0.01) was significantly higher at 7 days as compared to 3 days, supporting that there was cell growth under all conditions. C) Representative images of staining for F-actin (green) and nuclei (blue) in hASCs cultured dynamically or statically on DAT coatings or uncoated TCPS at 7 days (20% O_2_ condition shown). (n=2 replicate wells/trial, N=4 trials with different hASC donors). Scale bar represents 200 μm. Abbreviations: DAT, DAT coated; UC, uncoated TCPS.

We selected 2% O_2_ as the hypoxic condition to build on our previous work with the perfusion bioreactor system that showed positive effects of this level of hypoxia in combination with dynamic culture [28,29]. To select the rocking condition, a computational model of shear stress was developed using MATLAB, which was based on the work of Zhou *et al.* [31] with modifications to the modeling parameters to match the current rocking bioreactor system design. Based on the model, a rocking angle of ±11° and a rocking frequency of 50 rpm were selected to introduce the highest shear stress achievable in the system while keeping the cell culture fully immersed and avoiding splashing within the culture vessels (Supplementary Figure 1).

hASCs were seeded at a density of 5000 cells/cm^2^ on the DAT coatings or uncoated controls and cultured in complete DMEM/F12 proliferation media. The plates were incubated statically overnight under normoxic conditions to promote cell attachment (37 °C, 5% CO_2_). On day 0, the plates were moved to their assigned culture environment. The samples were then dynamically or statically cultured under 2% or 20% O_2_ for up to 7 days. For studies analyzing the effects of the culture microenvironment on hASC paracrine factor production using Human Magnetic Luminex^®^ assays or enzyme-linked immunosorbent assays (ELISA), the hASCs were cultured in complete proliferation media for 5 days and then transferred into low-serum media (DMEM/F12 supplemented with 2% FBS, 100 U/mL penicillin, and 0.1 mg/mL streptomycin) and cultured for an additional 48 h. For the later studies focused on analyzing the additional effects of priming with the pro-inflammatory cytokines TNF-α and IFN-γ on hASC paracrine factor secretion, hASCs were cultured in complete proliferation media for 5 days and then transferred into priming media (low-serum media supplemented with 10 ng/mL TNF-α and 100 ng/mL IFN-γ [32]) or low-serum media as controls and cultured for an additional 48 h.

### 2.4. Immunohistochemical analysis of hASC density

To evaluate the effects of the culture microenvironment on hASC growth, nuclear staining with Hoechst 33258 was used to quantify the cell density, in combination with phalloidin to visualize cell morphology. Samples (11°, 50 rpm: n=2 replicate wells/trial, N=4 trials with different hASC donors) were collected after 3 and 7 days of culture, fixed in 10% phosphate-buffered formalin for 10 min, permeabilized for 10 min (PBS, 0.1% Triton X-100), and blocked for 30 min (PBS, 0.1% Tween 20, 1% BSA). Next, the samples were stained with Hoechst 33258 (ThermoFisher, 1:1000) for 1 h at room temperature. Finally, the samples were mounted in Fluoroshield mounting medium (Abcam). Ten non-overlapping images were captured for each sample using an EVOS^®^ FL fluorescence imaging system (Thermo Fisher Scientific) under 20X magnification. Nuclei per mm^2^ were quantified using ImageJ software.

### 2.5 Intracellular IDO expression

To evaluate the effects of the culture microenvironment on intracellular IDO expression, hASCs were cultured as previously described (11°, 50 rpm, n=2 replicate wells/trial, N=3-4 trials with different hASC donors) and western blotting performed after 7 days of culture to quantify intracellular IDO production. Protein lysates were collected using RIPA lysis buffer with protein concentration quantified via the Pierce™ BCA Protein Assay Kit (Thermofisher) following the manufacturer’s instructions.

In brief, 10 μg of protein lysate from each group was combined with 4x Laemmli buffer (Bio-Rad) and heated to 95°C for 5 min. The Precision Plus Protein Dual Color Standards (Bio-Rad) and samples were then loaded into a 12% polyacrylamide gel created with the TGX Stain-Free™ FastCast™ Acrylamide Kit (Bio-Rad) following the manufacturer’s instructions. The gel was run at 200 V in running buffer (25 mM Tris, 190 mM glycine, 0.1% SDS) using the Mini-PROTEAN® Electrophoresis System (Bio-Rad). Proteins were then transferred to a low fluorescence polyvinylidene difluoride membrane using the Trans-Blot® Turbo™ Transfer System (2.5 A, 25 V, 10 min; Bio Rad) and Trans Blot® TurboTM RTA Transfer Kit (Bio-Rad).

The membranes were blocked (TBS +5% milk +0.1% Tween-20) for 1 h at RT under agitation at 200 rpm, followed by primary antibody staining against IDO (1:10,000 dilution, rabbit monoclonal, cat. Ab211017; Abcam) or the housekeeping protein TATA-binding protein (TBP) (1:1000 dilution, rabbit polyclonal, cat. Ab63766; Abcam) in TBS +5% BSA +0.1% Tween-20 overnight at 4°C, 100 rpm. Membranes were washed three times for 5 mins each in TBS +0.1% Tween-20 at RT, 200 rpm, and the secondary antibody (1:10,000 dilution, goat anti-rabbit HRP conjugated, cat. Ab6721; Abcam) was applied and incubated for 1 h in TBS +5% milk +0.1% Tween-20 at RT, 100 rpm. Membranes were then washed six times for 5 min each in TBS +0.1% Tween-20 at RT, 200 rpm. The membranes were imaged using the Clarity™ Western ECL Substrate System (Bio-Rad) on a ChemiDoc^TM^ MP Imaging System (Bio-Rad). Staining for TBP was completed first, and the membrane stripped using 100 mM 2-mercaptoethanol, 2% (w/v) SDS, 62.5 mM Tris-HCl, pH 6.7 in H_2_O, followed by blocking and staining for IDO. Band quantification was performed using ImageJ software with IDO expression normalized to TBP expression in each sample.

### 2.6 Luminex® analysis of pro-angiogenic and immunomodulatory factor secretion

To evaluate the effects of the culture microenvironment on hASC paracrine factor secretion, the ASCs were cultured as previously described (11°, 50 rpm: n=2 replicate wells/trial, N=3-4 trials with different ASC donors). On day 7, the conditioned medium (CM) from each sample was collected, centrifuged at 1200 × g for 10 min to remove particulates, and frozen at -80 °C. For the initial studies without cytokine priming, a 5-plex Luminex assay (angiogenin, VEGF, HGF, MCP-1, IL-8) was performed. For the subsequent studies evaluating the effects of cytokine priming, IL-8 and MCP-1 were analyzed in a duplex assay and angiogenin, VEGF, and HGF were combined in a triplex assay to accommodate the higher expression levels of the analytes in some samples. The Luminex Assays were performed in accordance with the manufacturer’s protocols using a MAGPIX^®^ System (Millipore). The results were analyzed using the xPONENT program with a five-parameter logistic curve-fit. Protein concentrations were determined based on comparison to the standard curves and normalized to the dsDNA content measured in each sample using a PicoGreen^®^ dsDNA Assay, described below.

### 2.7 ELISA analysis of IL-6 secretion

ELISA was performed to measure the levels of secreted IL-6 as it was much more highly expressed than the other secreted factors included in the Luminex panel. CM samples were processed and stored as previously described (n=2 replicate wells/trial, N=5 trials with different ASC donors). ELISA was performed in accordance with the manufacturer’s protocols using a CLARIOstar^®^ microplate reader (BMG Labtech, Guelph, Canada) at 450 nm and 540 nm. The results were analyzed using the MARS analysis program with a four-parameter logistic curve-fit. Protein concentrations were determined based on comparison to the standard curves and normalized to the dsDNA content measured in each sample using the PicoGreen^®^ dsDNA assay.

### 2.8 Picogreen^®^ dsDNA assay

PicoGreen^®^ dsDNA assays were performed following the manufacturer’s instructions, to normalize the secreted proteins levels in the CM samples based on the total number of cells in each corresponding well at 7 days. In brief, cell samples from the uncoated and DAT-coated conditions were collected using trypsin release or cell scraping, respectively. DNA extraction was performed using the DNeasy Blood & Tissue kit (Qiagen). Briefly, samples were digested overnight in 1 mL of buffer ATL supplemented with 32.5 μL of proteinase K. The dsDNA was then extracted from the digested samples and analyzed using the Quant-iT™ PicoGreen^®^ Assay kit (Invitrogen) according to the manufacturer’s instructions with a CLARIOstar^®^ microplate reader. The standard curve was generated using the Lambda DNA standard provided in the kit. Unseeded DAT coating samples were also included for background adjustment.

### 2.9 Statistical analyses

For analysis of ASC density and paracrine factor secretion, a mixed effects model was used to first evaluate the main effect of oxygen tension. The data were then divided into two groups based on the oxygen level and analyzed separately. To evaluate the effects of the culture microenvironment on ASC density and paracrine factor production, a two-way analysis of variance (ANOVA) test was performed, followed by a Tukey’s post-hoc multiple-comparisons test. To evaluate the effects of cytokine priming on ASC paracrine factor production, due to the additional variable (i.e., priming or non-priming media), a repeated-measures two-way ANOVA was performed, followed by a Tukey’s post-hoc multiple-comparisons test (comparing primed and non-primed under the same culture condition) and a Sidak’s post-hoc multiple-comparisons test (comparing between culture conditions under the same priming condition). To evaluate intracellular IDO content, a one-way ANOVA test was performed, followed by a Tukey’s post-hoc multiple-comparisons test. All statistical analyses were performed using the GraphPad Prism 8 software. All numerical values are represented as the mean ± standard deviation, and differences were considered statistically significant at p < 0.05.

## 3. Results

### 3.1 Effects of the culture conditions on hASC growth

The initial culture studies focused on applying the rocking bioreactor system to characterize the effects of combining the DAT coatings and shear-stress stimulation on hASC growth under 2% or 20% O_2_. Staining and quantification of cell nuclei was performed after 3 and 7 days of culture under the varying conditions using ImageJ. Under both 2% and 20% O_2_, the density of hASCs cultured on DAT coatings or TCPS was significantly higher at 7 days as compared to 3 days (Figure 2), supporting that there was cell growth in the system under all conditions. At day 3, the cell density was not significantly different between the culture conditions. However, at day 7, culturing under 20% O_2_ resulted in a significantly higher cell density as compared to under 2% O_2_ (p < 0.05). In addition, culturing on the DAT coatings under 20% O_2_ resulted in a significantly higher cell density at day 7 as compared to the uncoated group (p < 0.01) (Figure 2B). Staining of F-actin showed no obvious differences in hASC morphology induced by dynamic culture or culturing on the DAT coatings (Figure 2C).

**Figure 2.**
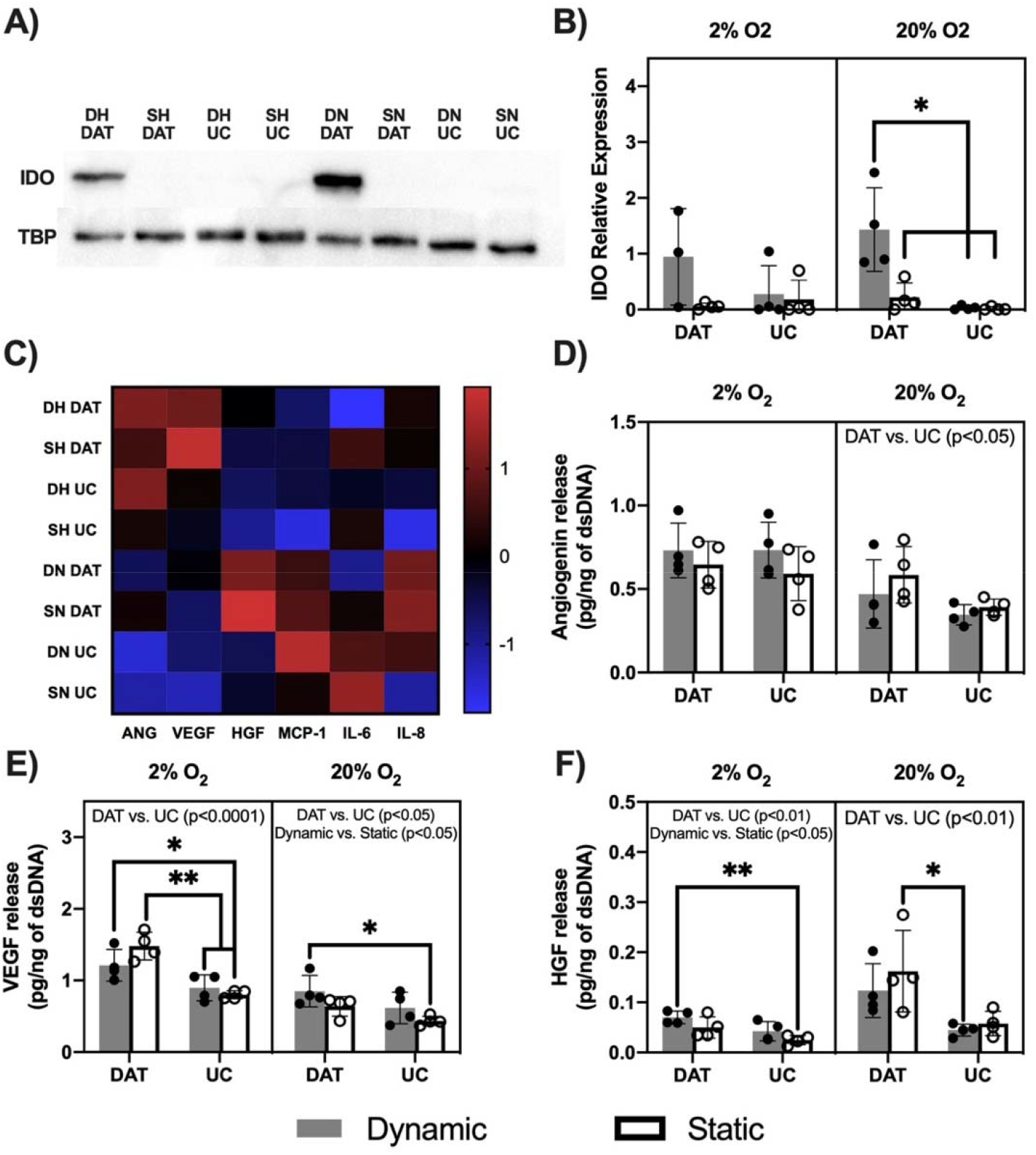
Culturing hASCs for 7 days under dynamic conditions and/or on the DAT coatings altered angiogenin, VEGF and HGF secretion. A) Representative western-blotting results of hASC lysates collected after 7 days of culture. B) Dynamic culture on DAT coatings enhanced intracellular IDO expression in hASCs cultured under 20% O_2_. C) Heatmap overview of the levels of secreted angiogenin, VEGF, HGF, MCP-1, IL-6 and IL-8 detected in the conditioned media samples. Data represents z-score average concentrations normalized to total dsDNA content. Secreted levels of D) angiogenin, E) VEGF, and F) HGF, normalized to total dsDNA content. Results of main effects analyses are noted above each plot. Post-hoc test results: *p < 0.05, **p < 0.01; n=2 replicate wells/trial, N=3-4 trials with different hASC donors. Abbreviations: ANG, angiogenin; D, dynamic; S, static; H, hypoxia; N, normoxia; DAT, DAT coated; UC, uncoated TCPS.

### 3.2 Pro-angiogenic and immunomodulatory paracrine factor secretion was modulated by the oxygen tension, dynamic culture, and DAT coatings

Following confirmation that all of the culture conditions supported hASC growth, the effects of combining the DAT coatings and shear-stress stimulation on intracellular expression levels of the immunomodulatory factor IDO were characterized through western-blotting analysis of cell lysates collected after 7 days of culture under 2% or 20% O_2_. Interestingly, visible bands for IDO expression were only observed in the hASCs that had been cultured dynamically on the DAT coatings (Figure 3A). Quantitative analysis confirmed that IDO expression was significantly enhanced in the hASCs cultured dynamically on the DAT coatings under 20% O_2_ (Figure 3B).

**Figure 3.**
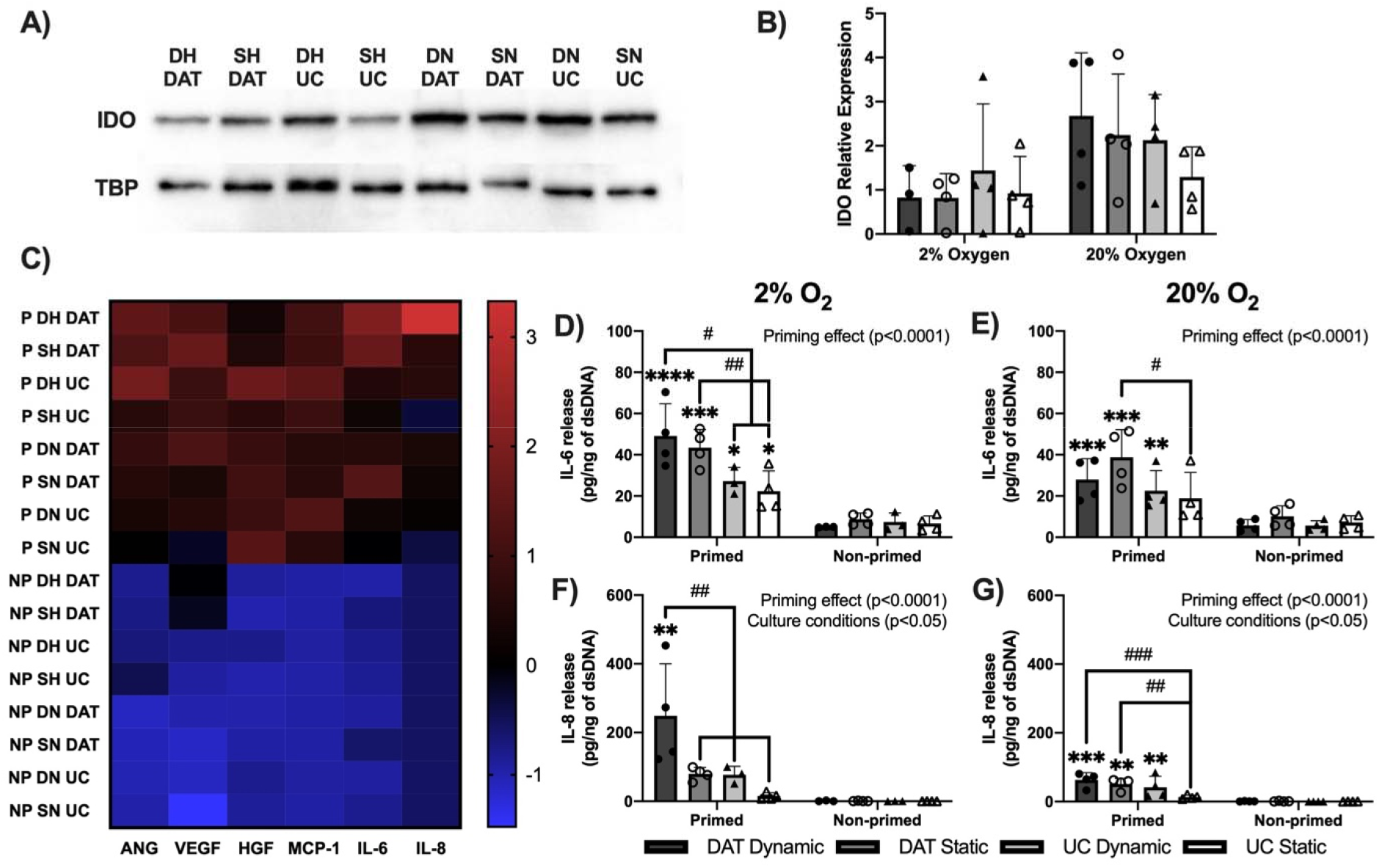
Priming with TNF-α and IFN-γ significantly enhanced the secretion of all tested paracrine factors in the human ASCs. A) Representative western blots of cell lysates collected after 7 days of culture (cytokine stimulation for the final 48 h) showing intracellular IDO expression in hASCs primed with TNF-α and IFN-γ. B) Quantification of western blotting showed no significant differences in the relative IDO expression levels between the primed groups. C) Heatmap overview of the levels of secreted angiogenin, VEGF, HGF, MCP-1, IL-6 and IL-8 detected in the conditioned media samples. Data represents z-score average concentrations normalized to total dsDNA content. Expression of IL-6 under D) 2% O_2_ and E) 20% O_2_ and IL-8 under F) 2% O_2_ and G) 20% O_2_, normalized to total dsDNA content. Results of main effects analyses are noted above each plot. Sidak’s multiple comparisons: *p < 0.05, **p < 0.01, ***p < 0.001, ****p < 0.0001 indicating significant difference between the primed and the non-primed sample under the same culture condition. Tukey’s multiple comparisons: #p < 0.05, ##p < 0.01, ###p < 0.001 indicating significant difference between the primed samples under different culture conditions. n=2 replicate wells/trial, N=3-4 trials with different ASC donors. Abbreviations: ANG, angiogenin; P, primed; NP, non-primed; D, dynamic; S, static; H, hypoxia; N, normoxia; DAT, DAT coated; UC, uncoated TCPS.

Next, CM samples were analyzed to assess the effects of combining the DAT coatings and shear-stress stimulation on the levels of secreted angiogenin, VEGF, HGF, MCP-1, IL-8, and IL-6 after a 7-day culture period under 2% or 20% O_2_, with media conditioning for the final 48 h. As reflected in a heat map of the results, the hASC paracrine factor secretion profile was modulated by the different microenvironmental conditions (Figure 3C). However, the expression levels of MCP-1, IL-6, and IL-8 secretion were not significantly altered by the varying culture conditions included in the current study (Supplementary Figure S2).

In terms of the main effects of oxygen tension, culturing under 2% O_2_ significantly enhanced the expression of VEGF (p < 0.05) and reduced the expression of HGF (p < 0.05) as compared to culturing under 20% O_2_. In terms of the main effects of the culture substrate, culturing on the DAT coatings significantly enhanced the expression of angiogenin under 20% O_2_ (p < 0.05, Figure 3D), as well as the expression of VEGF (2% O_2_: p < 0.0001, 20% O_2_: p < 0.05, Figure 3E) and HGF (p < 0.01, Figure 3F) under both oxygen tensions. In terms of the main effects of dynamic culture, culturing under dynamic conditions was shown to significantly enhance the expression of VEGF under 20% O_2_ (p < 0.05) and the expression of HGF under 2% O_2_ (p < 0.05).

Consistent with the main effects analysis, two-way ANOVA analyses indicated that VEGF secretion was significantly higher in the samples cultured dynamically on DAT coatings as compared to those cultured statically on TCPS (p < 0.05) under both 2% and 20% O_2_ (Figure 3E). Further, under 2% O_2_, VEGF secretion was significantly higher in the samples cultured statically on DAT coatings as compared to those cultured either statically or dynamically on TCPS (p < 0.01). In addition, HGF secretion was significantly higher in the samples cultured dynamically on DAT coatings as compared to those cultured statically on TCPS (p < 0.01) under 2% O_2_, (Figure 3F). In contrast, HGF secretion was significantly higher in samples cultured statically on DAT coatings as compared to those cultured dynamically on TCPS under 20% O_2_ (p < 0.05).

### 3.3 Priming with TNF-α and IFN-γ under dynamic culture significantly enhanced hASC secretion of IL-6 and IL-8

Recognizing that priming with pro-inflammatory cytokines can substantially augment hASC paracrine factor secretion, we sought to characterize the effects of stimulating the hASCs cultured within the rocking bioreactor with TNF-α and IFN-γ. Western-blotting analysis of IDO expression in cell lysates collected after 7 days of culture (cytokine stimulation for the final 48 h) showed visible bands in all cytokine primed groups, as expected (Figure 4A). However, there was cell-donor variability observed in the relative expression levels, with no statistically significant differences observed between the various primed groups (Figure 4B).

Characterization of the levels of angiogenin, VEGF, HGF, MCP-1, IL-6 and IL-8 detected in conditioned media samples revealed that cytokine priming had a dominant effect on the hASC secretion levels of the selected pro-angiogenic and immunomodulatory factors (Figure 4C).

Based on main effects analysis, the expression levels of angiogenin, VEGF, HGF, MCP-1, IL-6, and IL-8 were significantly higher in the primed samples as compared to the non-primed samples under both oxygen levels (Supplementary Figure S3). Under 2% O_2_ with cytokine priming, culturing dynamically on the DAT coatings significantly enhanced the expression of IL-6 as compared to the uncoated groups (p < 0.05) (Figure 4D). Similarly, culturing statically on the DAT coatings significantly enhanced the expression of IL-6 as compared to the statically cultured uncoated group (2% O_2_: p < 0.01, 20% O_2_: p < 0.05) (Figure 4E). Further, culturing dynamically on the DAT coatings under 2% O_2_ was shown to significantly augment the expression of IL-8 as compared to all other groups (p < 0.01) (Figure 4F). Under 20% O_2_ with cytokine priming, culturing on the DAT coatings statically (p < 0.01) or dynamically (p < 0.001) was shown to significantly increase the expression of IL-8 as compared to the statically cultured uncoated group (Figure 4G).

## 4. Discussion

A variety of preconditioning strategies have been explored to improve hASC expansion and paracrine factor production with the goal of enhancing their therapeutic efficacy for clinical applications [9]. Studies have focused on modulating a range of microenvironmental factors during culture including oxygen tension and the culture substrate, as well as stimulating cells mechanically or with growth factors and cytokines [33–37]. However, few studies have focused on assessing the combined effects of these various factors. To begin to address this gap, the current study developed a scalable rocking bioreactor system to enable the systematic characterization of the combined effects of DAT coatings, shear-stress stimulation, and pro-inflammatory cytokine priming on hASC proliferation and paracrine-factor secretion under varying oxygen tensions.

Importantly, staining results confirmed that hASC growth was observed under all of the culture conditions studied, supporting the potential of our rocking bioreactor system as a platform for hASC expansion. Notably, the DAT coatings were favorable for hASC growth under 20% O_2_, which is consistent with other studies reporting positive effects of DAT substrates on cell proliferation [4,19]. Moreover, the data indicated that hASC growth was enhanced under 20% O_2_, based on a higher cell density at day 7 compared to the samples cultured under 2% O_2_. Previous studies have reported that hypoxia can enhance hASC or MSC proliferation [38–41]. However, there are conflicting reports in the literature. For example, Frazier *et al.* reported no significant effects of a 72-h exposure to 5% O_2_ on hASC viability and proliferation [42]. Other studies have also suggested an inhibitory effect of hypoxia on ASC or MSC proliferation [43–45]. These differences highlight that many factors can influence the effects of hypoxia on cell proliferation, including the oxygen concentration in the medium, the duration of exposure, the specific cell source used, and the composition of the culture medium.

There has been relatively limited research to date on the application of fluid shear stress as a method of preconditioning cells. In cell-culture studies applying shear stress, the magnitude of the shear stress often varies substantially, ranging from hemodynamic fluid flow shear (e.g. 10 dyn/cm^2^ [23,24], 15 dyn/cm^2^ [25,26]) to low-level or interstitial fluid flow shear (0.5 dyn/cm^2^ [46,47], 0.018-0.024 dyn/cm^2^ [48]). These studies have primarily focused on unidirectional laminar flow [23,24,26,47,48], with shorter-term exposure to the dynamic culture conditions ranging from a few hours up to 4 days. To our knowledge, the current study is the first to explore the effects of longer-term exposure (up to 7 days) to low-level (0.04-0.3 dyn/cm^2^), non-uniform oscillatory fluid shear stress on hASC growth and paracrine-factor secretion.

While the shear stress applied in our rocking bioreactor system did not significantly impact the hASC density measured at day 7 in the current study, other studies have reported differences in proliferation patterns between shear-stress-stimulated and statically cultured ASCs. For example, Elashry *et al.* showed that a 10-day exposure to fluid shear stress (oscillatory, 0.77 dyn/cm^2^) increased proliferation of equine ASCs compared to static conditions [49]. Interestingly, another study by Kim *et al.* reported that the hASC proliferation rate was dependent on the level of shear stress applied [48]. More specifically, by comparing three regions in a microfluidic device with a shear-stress gradient, hASCs cultured in the lower shear-stress region (0.018 dyn/cm^2^) were found to have a higher proliferation rate compared to those in the higher shear-stress region (0.024 dyn/cm^2^).

To assess how culturing in the varying bioreactor-based microenvironments altered the paracrine profile of the hASCs, we first selected intracellular IDO as a functional marker of immunomodulatory and immunosuppressive properties of MSCs. IDO plays a key role in MSC-mediated immunosuppression by indirectly suppressing the T-cell response [50], inducing immunosuppressive regulatory T-cell (T_regs_) differentiation [51], and supporting monocyte differentiation into alternatively activated “M2-like” macrophages [27]. IDO is commonly known to be induced by IFN-γ or TNF-α and interleukin-1 beta (IL-1β) [27,35]. This is consistent with our results showing increased IDO expression among primed groups.

Interestingly, in the present study, we identified dynamic culturing combined with DAT coatings under normoxic conditions to be an additional regulator of the hASC intracellular IDO level. While IDO is strongly induced by inflammatory cytokines, particularly IFN-γ and TNF-α, the cellular microenvironment may also influence its regulation, although this remains poorly characterized. Zimmermann *et al*. reported that MSC aggregation into 3D spheroids did not significantly alter IDO activity compared to conventional 2D culture, suggesting that 3D culture alone is not enough to alter IDO activity [52]. In contrast, Duggal *et al.* demonstrated that MSCs encapsulated in RGD-functionalized alginate hydrogels exhibited increased expression of TDO2, a tryptophan-catabolizing enzyme related to IDO-mediated metabolic pathways, although regulated through distinct mechanisms [53]. In addition to structural cues, oxygen tension has been investigated as a regulatory factor. Consistent with our findings, Roemeling-van Rhijn *et al*. and Wobma *et al*. found that hypoxia alone did not significantly alter IDO expression or activity relative to normoxic conditions [54,55]; however, Wobma et al. reported a synergistic effect of hypoxia with IFN-γ priming on IDO activity compared to priming alone [55]. To the best of our knowledge, this is the first study to assess the effects of dynamic culture on hASC intracellular IDO expression.

Next, Luminex and ELISA assays were performed to evaluate the levels of angiogenin, VEGF, HGF, MCP-1, IL-6, and IL-8 detected in conditioned media samples collected after a 7-day culture period, with media conditioning for the final 48 h. Comparing between the DAT-coated group and the uncoated TCPS group, the pro-angiogenic factors VEGF, HGF, and angiogenin (20% O_2_ only) were detected at significantly higher levels in the DAT-coated groups. These findings are consistent with our previous data demonstrating enhanced angiogenic-factor secretion when human fibroblasts where cultured on 3D DAT bioscaffolds as compared to structurally similar collagen scaffolds [56]. Taken together, these data support that DAT can function as a cell-instructive platform to augment the pro-angiogenic potential of therapeutic cell populations.

In terms of the effects of shear-stress stimulation, the current study demonstrated that VEGF (20% O_2_) and HGF (2% O_2_) secretion were upregulated in the dynamically cultured groups. Similarly, Bassaneze *et al.* demonstrated a shear-stress induced nitric oxide (NO)-dependent VEGF accumulation in hASC conditioned media, with an increasing production rate over a 96-h shear-stress exposure (unidirectional, 10 dyn/cm^2^) [24]. In addition, Bravo *et al.* reported increased secretion of VEGF and HGF – along with other pro-angiogenic factors – when hASCs were exposed to 10 min of shear stress (unidirectional and intermittent, 10 dyn/cm^2^) as compared to statically cultured controls [23]. Providing evidence that the hASC secretome has functional effects relevant to vascular regeneration, Chen *et al.* reported improved viability of human endothelial cells in response to oxidative stress when cultured with conditioned media collected from shear-stress-stimulated hASCs (unidirectional, 0.5 dyn/cm^2^, 30 min) as compared to statically cultured hASCs [47].

In the current study, considering the combined effects of shear stress and hypoxia, an interesting secretion pattern of VEGF can be observed that is somewhat aligned with the Bravo *et al.* study, which revealed that shear stress and hypoxia had opposite effects on hASC pro-angiogenic factor secretion [23]. More specifically, it was reported that shear stress and hypoxia each independently enhanced hASC pro-angiogenic capacities via cyclooxygenase-2 (COX-2)-dependent mechanism. However, when shear stress and hypoxia were combined, VEGF secretion was significantly reduced compared to introducing hypoxia alone. This result may partially explain why dynamic culture only significantly enhanced VEGF secretion under 20% O_2_ in the current study.

Pro-inflammatory cytokine priming is another widely investigated preconditioning strategy to enhance the immunomodulatory function of MSCs [35]. Under IFN-γ and TNF-α priming, in addition to upregulation of IDO [27], it has been reported that MSCs secrete higher levels of VEGF, HGF, IL-6, and IL-8 [35,57,58]. In the current study, the expression levels of all tested paracrine factors were significantly upregulated in the primed groups based on main effects analysis. Interestingly, the immunomodulatory factors MCP-1, IL-6 and IL-8 were upregulated far more by priming than the angiogenic factors angiogenin, VEGF, and HGF. This phenomenon has not been explicitly reported in the literature, which warrants further investigation of the mechanism behind the upregulation of these paracrine factors under the effect of priming.

To the best of our knowledge, this is the first study to evaluate hASC paracrine factor secretion under different culture conditions within a bioreactor system in combination with IFN-γ and TNF-α priming. Under the priming conditions, the immunomodulatory factors IL-6 and IL-8 were most impacted by the varying culture conditions (DAT coating, shear-stress stimulation) within the bioreactor. Interestingly, in contrast, the pro-angiogenic factors VEGF, HGF, and angiogenin were most affected by the culture conditions without priming. Although generally regarded as a pro-inflammatory cytokine, IL-6 is involved in many aspects of the immunoregulatory activities of MSCs, including promoting the polarization of macrophages towards an alternatively activated M2-like phenotype [59,60], inhibiting monocyte differentiation towards dendritic cells [61], and supporting the production of prostaglandin E2 (PGE2) and NO [62]. IL-8 on the other hand plays a key role in MSC-mediated endothelial progenitor cell homing and angiogenesis [63]. As such, future studies are warranted to investigate whether the increased levels of IL-6 and IL-8 secreted by the preconditioned hASCs could be beneficial from a therapeutic perspective.

Future studies should also evaluate the effects of CM generated by hASCs using more functional assays. For example, hASC pro-angiogenic function could be assessed by characterizing the effects of CM on human microvascular endothelial cell proliferation, viability, and tubule formation [64,65]. Additionally, endothelial barrier function could be characterized using a vascular permeability assay kit, as vessel stabilization by the hASCs may be an important mechanism of regeneration [66]. Additionally, the immunomodulatory capacity of hASCs could be tested by applying CM to peripheral blood-derived monocytes and assessing differentiation and polarization via flow cytometry. Similarly, the effects of CM on macrophage function can be assessed *in vitro* using a series of bead-based phagocytosis and efferocytosis activity assays [67] as well as the chemotactic effects of CM through a macrophage transwell migration assay [68,69]. Depending on the outcomes of the *in vitro* functional assays, future studies could also characterize the effects of hASC preconditioning on *in vivo* angiogenesis using preclinical rodent models [70].

## 5. Conclusions

In conclusion, a novel bioreactor system was established for hASC expansion and preconditioning, and the combined effects of DAT coatings, shear-stress stimulation, and varying oxygen tensions were assessed. The bioreactor supported the expansion of hASCs under all culture conditions, with enhanced intracellular IDO expression in the hASCs dynamically cultured on DAT coatings under 20% O_2_. Notably, shear-stress stimulation and/or culture on DAT coatings also enhanced the secretion of the pro-angiogenic factors VEGF, HGF, and angiogenin. When hASCs were primed with the pro-inflammatory cytokines TNF-α and IFN-γ, intracellular IDO expression was detected in all groups, and the levels of all secreted factors detected in the conditioned media samples were enhanced, particularly IL-6, IL-8, and MCP-1. Interestingly, under cytokine priming, IL-6 and IL-8 expression was most affected by the varying microenvironmental factors studied (i.e. shear-stress stimulation & DAT coatings). Overall, this study demonstrated that the cell microenvironment can be tuned to modulate hASC paracrine factor secretion and supports the further investigation of bioreactor platforms integrating shear-stress stimulation and ECM-derived biomaterials as a strategy for cellular preconditioning to leverage the hASC secretome for regenerative applications.

## Data Availability Statement

The raw data supporting the conclusions of this manuscript will be made available by the authors, without undue reservation, to any qualified researcher.

## Acknowledgements

This study was supported by operating grant funding from the Canadian Institutes of Health Research (CIHR Institute of Musculoskeletal Health and Arthritis Priority Announcement #179858) and Natural Sciences Research Council of Canada (NSERC RGPIN-2017-0410). Zhiyu Liang was supported by a Transdisciplinary Bone & Joint Training Award from the Bone and Joint Institute at Western University, and Connor Gillis was supported through Ontario Graduate Scholarship (OGS) and NSERC Canada Graduate Scholarship Awards. The authors thank Drs. A. Grant and D. Matic for their clinical collaborations with the provision of adipose tissue samples, as well as Dr. Eric Arts for providing access to the Bio-Plex® MAGPIX™ Multiplex Reader system for the Luminex® analyses.

