## Supplementary Figures for "Development of a rocking bioreactor strategy to augment pro-angiogenic factor secretion by human adipose-derived stromal cells"

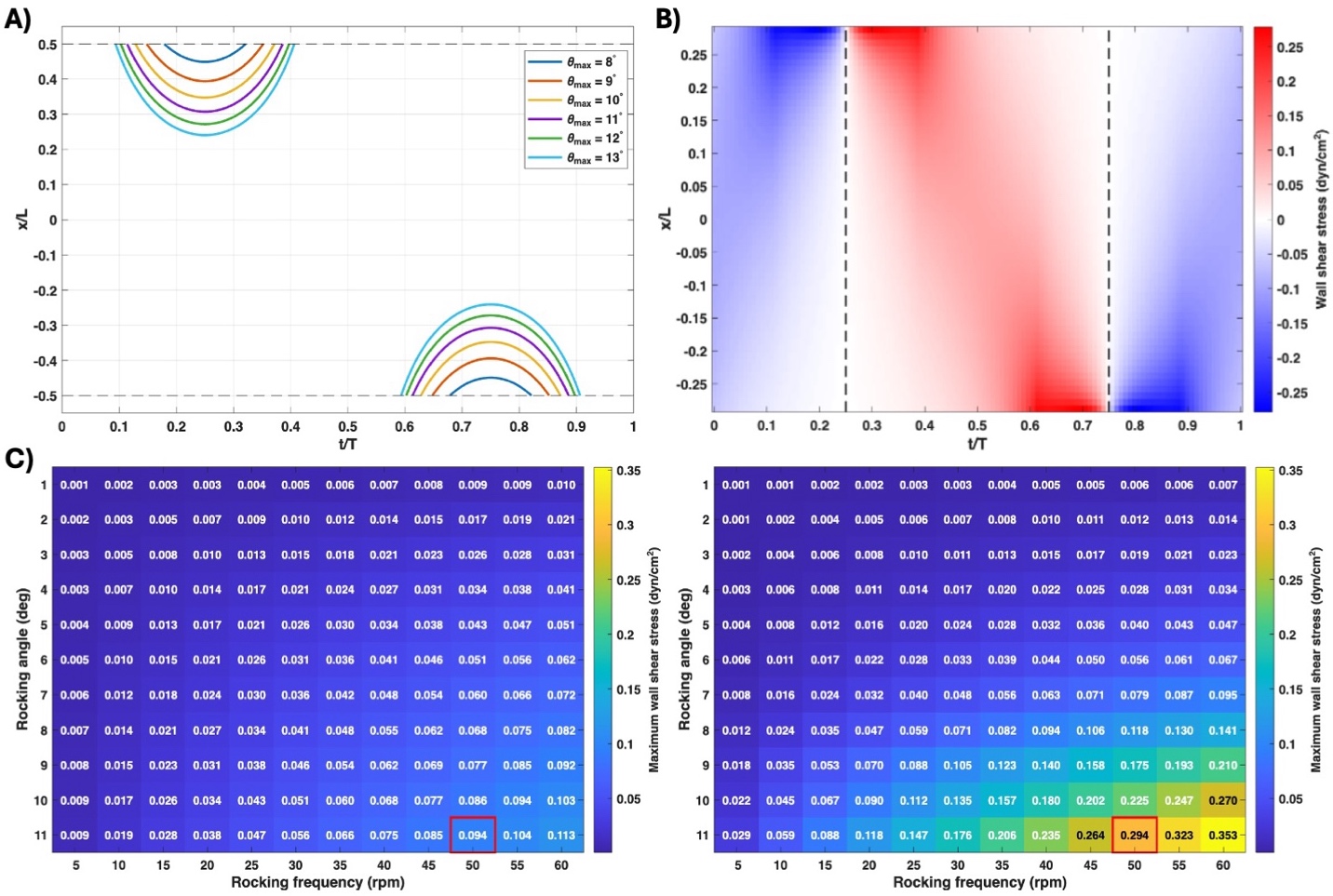

**Supplementary Figure S1. Computational model of shear stress simulation within the rocking bioreactor system.** A computational model of shear stress was developed in MATLAB, adapted from the previous work of Zhou et al. [31]. A) A simulation of the movement of 2.5 mL of fluid (height 2.38 mm) within an individual 27.9 mm x 37.6 mm culture chamber as a function of time and rocking angle demonstrates the boundary of the air-fluid interface on the floor of the culture chamber. Selecting a rocking angle of 11° or less ensured that a coverslip of 22-mm length at the centre of the chamber remained fully covered by the fluid throughout the rocking cycle. X axis: relative location (x/L) based on position x along the length of the well (L), Y axis: relative time (t/T) over a single rocking cycle (T). B) Simulation of the fluid shear stress (dyn/cm^2^) at the surface of the cell culture as a function of relative position and time for the selected rocking angle of 11° and rocking frequency of 50 rpm. The vertical dashed lines indicate the time of maximum tilt angle followed by change in flow direction. C) Maximum fluid shear stress under different rocking conditions shown at the centre of the well (x = 0) (left panel) and at the edge of the coverslip in the well (x = ±11 mm; x/L = ± 0.29) (right panel) as a function of rocking angle and frequency. The colour gradient from blue to yellow shows an increase in the value of the maximum shear stress, which increases at both locations as the rocking angle and frequency increase. The selected rocking condition of 11° and 50 rpm is indicated in the red box, corresponding to panel (B) above.

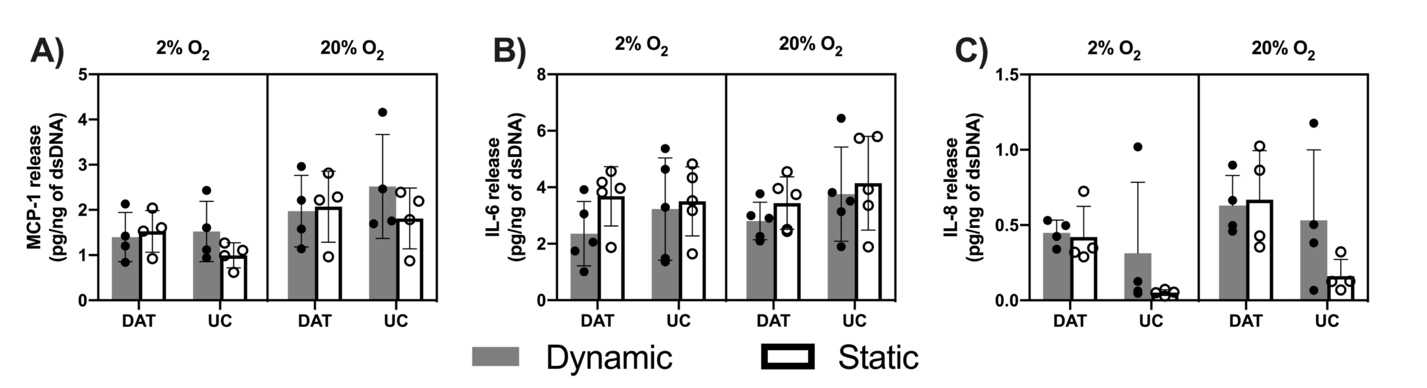

**Supplementary Figure S2. Culturing human ASCs for 7 days under dynamic conditions and/or on the DAT coatings did not significantly alter MCP-1, IL-6, and IL-8 secretion.**

Expression of A) MCP-1, B) IL-6, C) IL-8, normalized to total dsDNA content. n=2 replicate wells/trial, N=4 trials with different ASC donors, except IL-6: n=2 replicate wells/trial, N=5 trials with different ASC donors). Abbreviations: DAT, DAT coated; UC, uncoated TCPS.

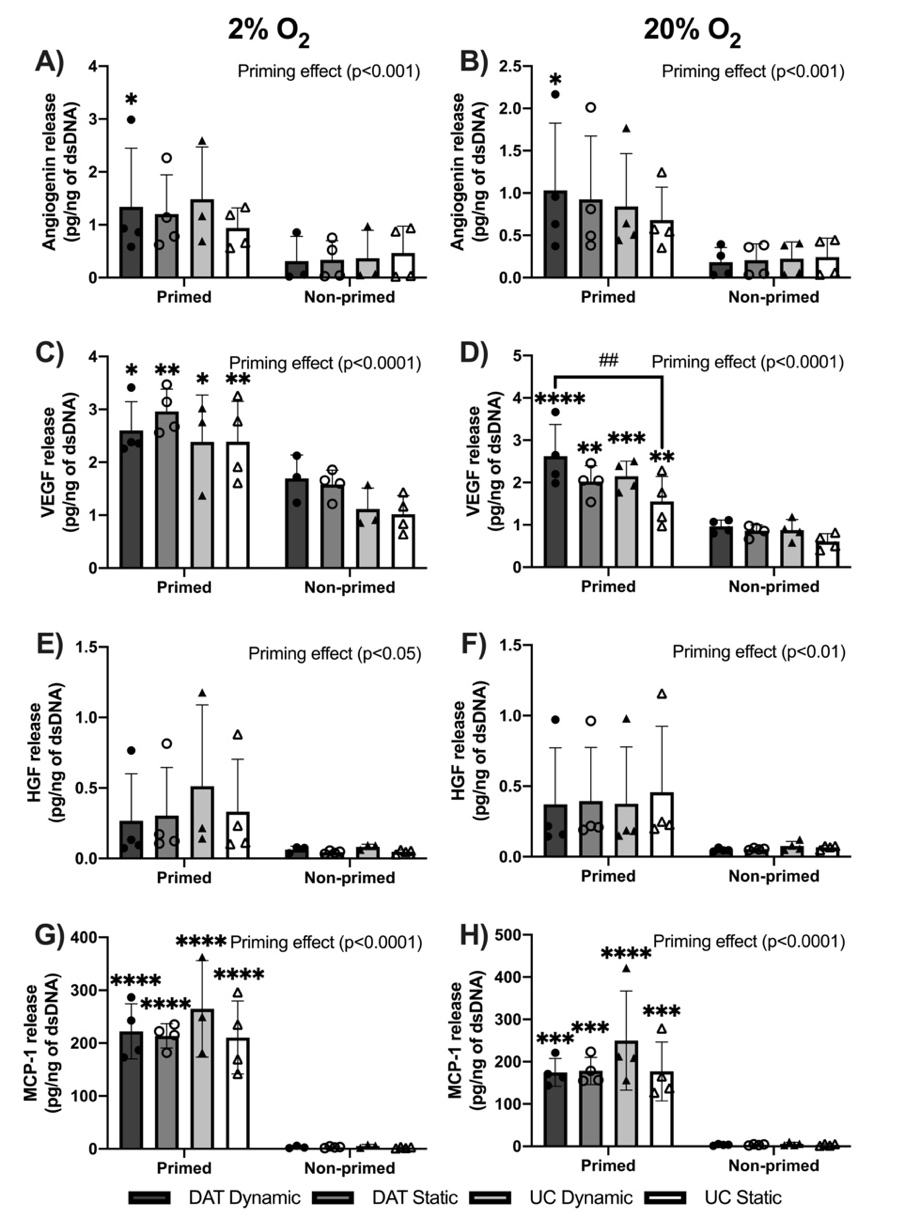

**Supplementary Figure S3. Priming with TNF-α and IFN-γ significantly enhanced the secretion of angiogenin, VEGF, HGF, and MCP-1 in the human ASCs.** Levels of secreted angiogenin under A) 2% O_2_ and B) 20% O_2_, VEGF under C) 2% O_2_ and D) 20% O_2_, HGF under E) 2% O_2_ and F) 20% O_2_, and MCP-1 under G) 2% O_2_ and H) 20% O_2_, normalized to total dsDNA content. Cytokine priming, as a main effect, significantly enhanced the expression of angiogenin (p < 0.001), VEGF (p < 0.0001), HGF (2% O_2_: p < 0.05, 20% O_2_: p < 0.01), and MCP-1 (p < 0.0001). Sidak’s multiple comparisons: *p < 0.05, **p < 0.01, ***p < 0.001, ****p < 0.0001 indicating significant difference between the primed and the non-primed sample under the same culture condition. Tukey’s multiple comparisons: ##p < 0.01 indicating significant difference between the primed samples under different culture conditions. n=2 replicate wells/trial, N=3-4 trials with different ASC donors. Abbreviations: DAT, DAT coated; UC, uncoated TCPS; P, primed; NP, non-primed.
